# Discerning state estimation and sensory gating, two presumptive predictive signals in mouse barrel cortex

**DOI:** 10.1101/2023.12.23.573180

**Authors:** Kalpana Gupta, Ritu Roy Chowdhury, Shubhodeep Chakrabarti, Cornelius Schwarz

## Abstract

The brain is assumed to contain distinctive predictive systems ranging from sensorimotor to cognitive functions. Here we report the successful functional delineation of two different presumptive predictive systems on the neuronal level, state estimation (SE) and sensory gating (SG), which both attenuate sensory flow during movement. Studying neuronal sensorimotor responses throughout the depth of primary somatosensory cortex in mice trained on a whisker reach task, SE appeared as a learned attenuation of tactile signal flow, due to a suppressive predictive signal, precisely at the time of an experimental sensory consequence. In contrast, SG was observable during a much longer interval after onset of the motor command. Both phenomena are internal, presumptively predictive signals, as blockade of the reafference did not affect them. We speculate that SG may be related to cognitive signals monitoring goals of movements, while SE likely is the expression of the classical notion of the reafference principle.

## Introduction

Generating predictions is one of the most important brain functions. They come at different hierarchical levels, from sensorimotor to cognitive, and likely engage an overlapping set of computationally powerful brain structures. The aim of this study was to delineate two well-studied phenomena, both likely related to predictive functions, called state estimation (SE) and sensory gating (SG). Both processes attenuate peripheral sensory signals in a movement dependent fashion, a commonality that, so far, was the reason for some uncertainty about whether they might or might not implement different functions (SG: ^1–4^; SE: ^5,6^). SE likely is the expression of prediction of sensory consequences of self-motion, the reafference signal ^6–8^. It is specific to movement kinematics and predicting sensory consequences at high temporal precision, and is thought to involve the cerebellum or cerebellum-like structures ^5,9–11^. The comparison of the predicted sensory consequences with the actually arriving sensory feedbacks, calculating a sensory prediction error, have been suggested to involve also the primary somatosensory cortex (S1)^12^. On the other hand, SG’s function is far more obscure. Its attenuation occurs along the entire ascending tactile pathway ^13,14^, and is independent of detailed movement kinematics and timing, thus seemingly independent of reafferent signals, and simply triggered by self-movement ^2,15^. SG-attenuation is dependent on intactness of S1 and surrounding parietal cortex ^13^. These results suggest that S1 is a site playing key roles in both presumptive predictive systems, where their functional separability can be studied (Fig. 1A). Psychophysical performance of humans confronted with a combination of force matching and sensory gating tasks, recently gave evidence for separability of two similar systems, studied on the behavioral level ^16^.

**Figure 1:**
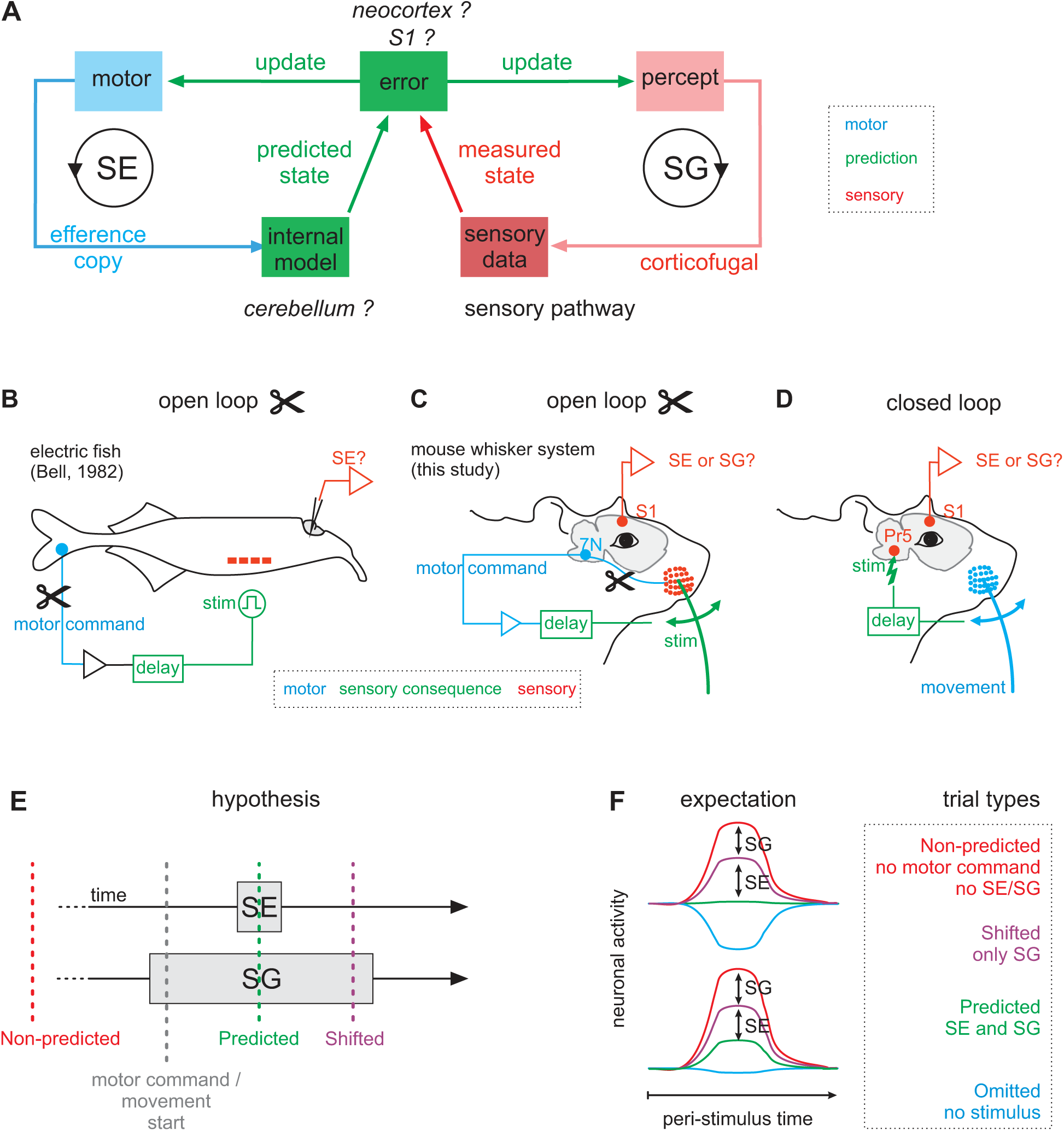
Hypothesis and experimental strategy. **A:** Hypothetical functional circuit model generating SG (right side) and SE (left) (italics: hypothesized brain structures). SE is the modern formulation of the reafference principle^7^. An internal model (green) receives the motor command (efference copy, blue), to calculate predictions about sensory consequences of a movement. The predicted state is then compared to the actual sensory feedback (red) to generate a sensory prediction error ^6^. SG (function unknown) acts on the ascending sensory pathway in a top-down fashion^13^ (pink). **B:** Bell’s open loop approach in weakly electric fish ^10^. The loop was opened by curarizing the electromotor cells, recording the motor command on the skin, and substitute the natural consequences of an electric organ discharge by an electrical stimulus in the fish tank. The predictive signals were recorded in the lateral line lobe, a cerebellum-like structure. **C:** In this study, we paralyzed whisker movement by severing the facial nerve, recorded the motor command upstream of the lesion in brainstem facial nucleus motoneurons, and triggered precise whisker deflection directed to the paralyzed whisker using an actuator, as a substitute sensory consequence. Predictive signals were assessed from multi-neuron recordings in S1 (barrel cortex). **D:** Closed loop approach. Whisker movements are intact, and sensory consequences are presented via microstimulation of the ascending whisker-related tactile pathway. **E:** Logic to delineate SG and SE. We posited that SG is active during the entire movement (large box), while SE acts in temporally precise fashion around the learned delay of the stimulation (small box). We tested the response of S1 presenting 4 different stimulus conditions (one per trial): ‘Predicted’, ‘Shifted’, ‘Omitted’, and ‘Non-predicted’. Predicted (green, 77% of trials) presented a whisker flick at the learned (and therefore predicted) delay after onset of motor command. Shifted (violet, 7.7% of trials) was shifted in time to tap into SG, but not into SE. Non-predicted (red, 7.7% of trials) trials presented a whisker flick in the absence of a motor command. Finally, Omitted (blue, 7.7% of trials) presented no stimulus at all. **F:** Expected attenuation of S1 tactile signals with the four trial types. Red: full tactile response (Non-predicted trials). Violet: Medium attenuation because Shifted trials would engage only the neuronal mechanism underpinning SG. Green: Maximal attenuation because Predicted would engage both systems. Blue: Omitted would reveal the internal prediction signal. Effect of SG: difference in response between Non-predicted and Shifted (red – violet). Effect of SE: difference of Shifted and Predicted (violet – green). Two possible outcomes are shown. Top: SE and SG together would abolish the tactile signal, and Omitted would reveal a suppressive signal. Bottom, incomplete attenuation (green), potentially obscuring the prediction signal (blue).

The study of predictive systems is based on some type of perturbation of sensory consequences of movement, an experimental interference, which is sometimes referred to as “opening the sensorimotor loop”. However, the experiments reported here go beyond perturbation, as in addition, they use anatomical means to manipulate the movement-related tactile feedback (i.e. the reafference). This strategy helps to argue that the origin of observed movement dependent modulation is generated by central signals rather than simply being due to peripheral sensory feedback ^15^. The critical element of our approach is to open the sensorimotor loop by cutting the motor nerve, and thus, block movement-related sensory feedback. We developed a real-time brain machine interface to assess the intact motor command proximal to the lesion and to trigger artificial sensory consequences and record their reflections in the brain. Figure 1BC reveals that our open loop paradigm in mice directly relates to the classic experimental approach of this type in the weakly electric fish^10^. For the purposes of the present study we will only refer to the experiments with anatomical blockade as “open loop” experiments. As a control and reference, however, we will compare the results gained in the open loop situation to results obtained with intact reafferent signaling, which we call the “closed loop” approach (Fig. 1D). The closed loop approach has been most commonly used in the literature (e.g. ^1,2,7,8,11,17–21^). It is therefore worth noticing that our terms “open” and “closed loop” refer to anatomical blockade, not to stimulus manipulations.

The logical basis to separate the two presumptive prediction systems is our working hypothesis that SE-attenuation is temporally precise (to be able to eliminate sensory consequences of movement), while the SG internal prediction is dispersed over rather long stretches of the movement. To test this hypothesis, we measured barrel cortex responses (whisker-related primary somatosensory cortex in rodents) to the learned and predicted artificial sensory consequence (a whisker flick, green in Fig. 1E, tagged ‘Predicted’) and compared these responses to test trials where we shifted the consequence to other latencies than the predicted ones (violet in Fig. 1E, tagged ‘Shifted’). We also placed test trials outside the movement period (red, tagged Non-predicted) to measure the total sensory response that can be elicited. Finally, omission of the predicted stimuli should reveal the predictive signal (tagged ‘Omitted). Figure 1F shows two scenarios of expected stimulus evoked tactile responses under these assumptions (top: incomplete; bottom: complete attenuation). The results reported here gave strong support for this hypothesis.

## Results

The experiments of the present study are based on an established whisker reach task in head-fixed mice ^22^. Before behavioral training, the mice were implanted with a single recording electrode in facial nucleus (7N) at a site where microstimulation evoked whisker movements, and either a 2×2 mobile wire-electrode-array (7 mice), or a 16-electrode linear silicon-electrode-array (5 mice), in the C2 barrel column. Mice were trained to retract their whisker to a starting position behind a baseline. Once ready, they self-initiated a rapid whisker reach across the baseline and a target located rostral to the baseline. Both, baseline and target were virtual computer-controlled whisker positions, the whisker moved across them without touching any real object. Reaching the target, the mice received a water reward (Fig. 2A-C). A trial was aborted if the animal issued a lick within an interval of 200 ms after reaching the target, or if the whisker did not move across baseline and target in an interval of 50 ms. Figure 2D-F exemplifies characteristic variable distributions of whisker reaches (duration and velocity, Fig. 2DE), latency of a 7N motor command to a movement (Fig. 2F).

**Figure 2:**
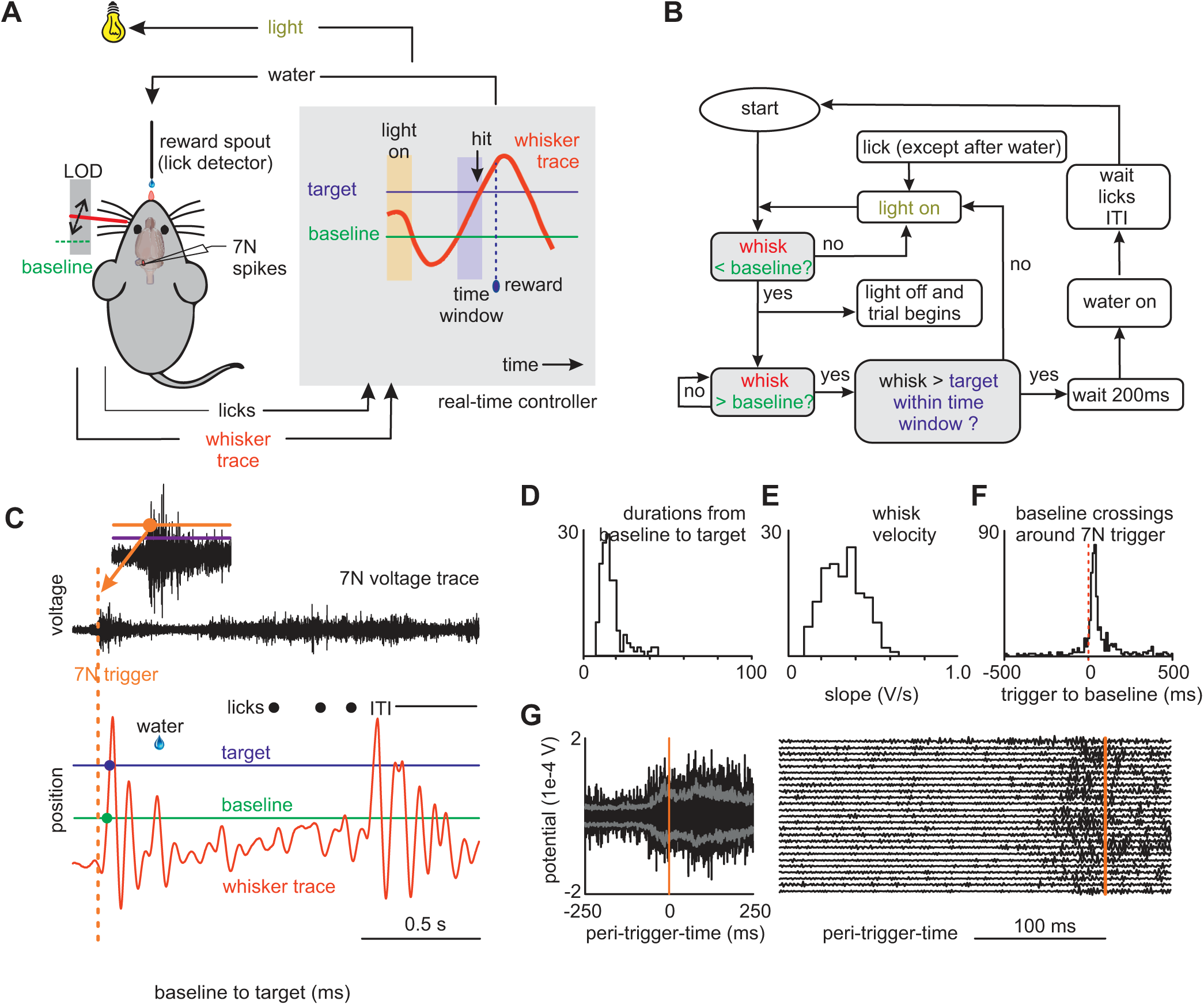
Training of a whisker reach task. **A:** A real-time controller registers licks as well as whisker movement of a head-fixed trained mouse. The mouse needs to bring the whisker behind a baseline (green) for 100 ms before a trial can be started. The trial is started by moving the whisker across the baseline (green). The reaching of a target maximally 50 ms later is required to count as a successful trial and is rewarded by delivering a drop of water. The house light is used to signal the whisker position to the mouse (before [ON] and behind the baseline [OFF]). LOD: laser-optical device to track whisker movement. **B:** Flow diagram of the state engine realizing automatic experimental control. **C:** One successful trial. Whisker position trace (red), baseline (green, dot: whisker crossing time), target (blue, dot: whisker crossing time). The recording in 7N is shown on top. At this stage, the 7N activity has no control function, but it is recorded to set thresholds needed to determine an 7N motor command (violet: used to determine absence of activity in 7N; orange: to determine the onset of a motor command [orange dot]). **D-F:** Whisker reach parameters. Examples of a typical mouse. Distribution of duration of whisk from baseline crossing to target crossing (D), distribution of maximum whisker velocity (E), baseline crossings following 7N triggers (F). **G:** Left: Example of 7N recordings triggered to 7N triggers (time zero) (776 recordings plotted on top of each other; gray lines: 25 and 75 percentile). Right: a subset of these recordings plotted out one by one. Orange vertical lines: 7N trigger time.

After successful training of the whisker-reach, we conducted recording sessions, in which we established the correspondence of multi-unit activity in 7N and whisker movement. 7N activity during a whisker reach movement showed an abrupt onset as reported earlier ^23^ (Fig. 2C). To count as a valid 7N trigger event we set the condition that 7N activity was absent for minimally 500 ms (assured by requiring 7N activity below a baseline, violet line in Fig. 2C), followed by an abrupt burst of activity (as detected by surpassing a threshold, orange in Fig. 2C). Figure 2G demonstrates 7N activity with respect to 7N trigger time.

### Open loop experiments

Next the mice were transferred to the open loop experiment. To paralyze whisking on the left side of the snout we severed the ipsilateral marginal mandibular as well as the buccal branches of the facial nerve, following Semba and Egger ^24^ (the sixth entry in their table 1). As reported in rats, these cuts completely paralyzed ipsilateral whisker movements also in the 12 mice used in this study. The mice were then trained on the behavioral task, but now triggering the start of a trial on 7N burst activity, i.e. a whisker motor command, as characterized during whisker reach training before nerve lesion (Fig. 3A-C). A brain-machine interface was established based on a real-time state engine to monitor 7N activity with respect to baseline and threshold of neuronal multi-unit activity at a temporal resolution of 0.1 ms. To enable a trial, the mice had to keep 7N activity below a baseline for a minimum interval of 500 ms, upon which the light would go off to signal to the mice that the trial start is now enabled ^22^. After the animals initiated the motor command, the trial started and a sensory consequence would be presented with a pre-defined delay with respect to its onset. The sensory consequence consisted in a short pulsatile deflection of the immobilized whisker of 10° amplitude and 10 ms duration using an actuator. One such deflection would be presented per trial (in some trials the consequence was omitted, i.e. there was no deflection at all). After licking off the water reward an intertrial interval would ensue and a new trial could be enabled by moving the whisker behind the baseline.

**Figure 3:**
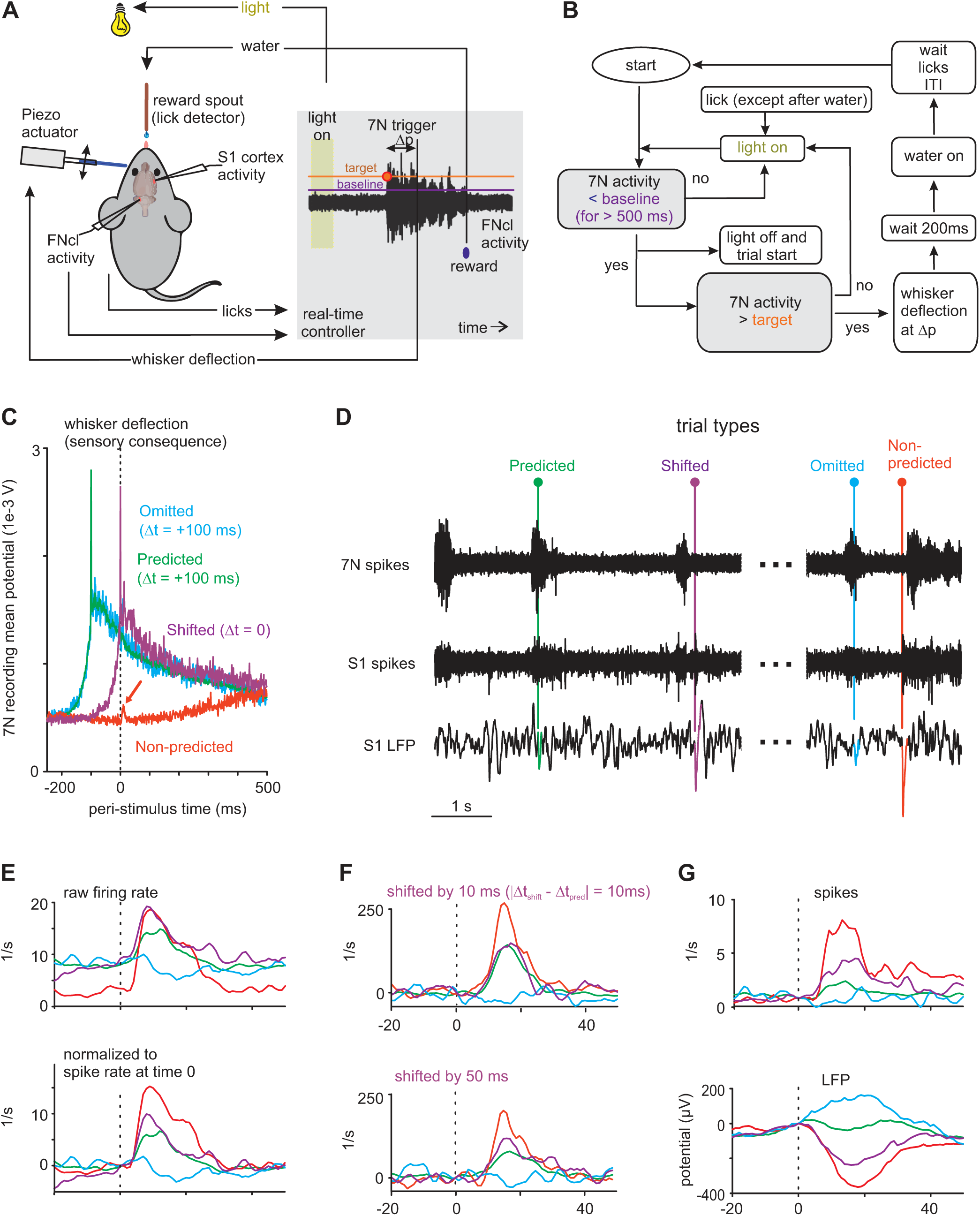
Open loop experiment: Training of 7N motor command. **A:** A real-time controller registers licks as well as the 7N recording. The 7N activity is required to stay below a baseline (violet, adjusted individually for each mouse) for 500 ms before a trial can be started by 7N activation exceeding a target (orange). A successful trial is rewarded by delivering a drop of water. The house light is used to signal the 7N to the mouse (OFF / ON signals below / above the baseline). **B:** Flow diagram of the state engine realizing automatic experimental control. **C:** 7N activity (typical example, mean across trials of one session). The red arrow indicates a sensory response in the 7N motor command, inconsistently seen most prominently in Non-predicted responses (in which motor signals are absent). **D:** Example raw voltage data. The automatic experimental control determines the sequence of trials, i.e. the type of stimulation (color). On top 7N activity triggering the stimuli. Center and bottom S1 spike and LFP recordings. The LFP responses to different stimuli types have been colored respectively. The recording exemplifies the ranking of response amplitudes found in this study (red→violet→green→blue = Non-predicted→Shifted→Predicted→Omitted). **E-G:** Example responses from different well-trained animals. Averages across one session. **E:** Top: pre-stimulus spike rates as recorded. Bottom: same data but all rates normalized to zero at time zero. **F:** Top: Responses obtained from a session that presented Predicted and Shifted at a difference latency of 10 ms. Bottom: Same animal different session with latency difference 50 ms. **G:** An example of spike rates (top) and LFP (bottom), in which the maximal attenuation (green) was almost total. Delays of stimuli presentations with respect to motor command onset: Predicted: 100ms (all panels), Shifted: 0 (C), 500 ms (D), 200 ms (E,G), 110 ms and 150 ms (F).

Table 1.

**Table 1.**
| Experiment | Delay conditions |  | Spikes |  |  | LFPs |  |  |
| --- | --- | --- | --- | --- | --- | --- | --- | --- |
|  | Predicted Delay (ms) | Shifted Delay (ms) | No. of animals | No. of delay conditions (electrodes) | No. of sessions | No. of animals | No. of delay conditions (electrodes) | No. of sessions |
| Ranking Open loop Fig. 4 | 100 | 0/300/400/500/ random | 12 | 80 (42) | 176 | 10 | 83 (40) | 157 |
| Ranking Closed loop Fig. 10 | 0, 100 | 50/62.5/75/100/200/250/300/350/375 | 4 | 46 (11) | 45 | 4 | 30 (8) | 27 |
| SE Precision Fig. 6A-C | 100 | 110 | 5 | 16 | 19 | 5 | 17 | 19 |
|  | 100 | 150 | 5 | 16 | 26 | 5 | 17 | 25 |
|  | 100 | 200 | 5 | 16 | 26 | 5 | 17 | 25 |
| SE range Fig. 7A-C | 0 | 100/250/300/500 | 4 | 9 | 46 | 3 | 11 | 36 |
|  | 200 | 300/400/ random | 6 | 23 | 43 | 6 | 30 | 40 |
|  | 400 | 0/random | 4 | 20 | 24 | 4 | 24 | 24 |
| Learning Fig. 8 | 100 | 300/400 | 4 | 15 | 4 | 3 | 14 | 4 |
|  | 200 | 300/400 | 4 | 15 | 4 | 3 | 14 | 4 |
| SG range Fig. 9A-C | 100 | 500 | 6 | 19 | 36 | 6 | 20 | 36 |
|  | 100 | 700/1000 | 4 | 21 | 16 | 4 | 23 | 16 |

Total Sample. Ranking of responses to Non-predicted, Shifted, Predicted, Omitted. The gray columns hold the sample n used for statistics

Before recording commenced, a few behavioral sessions were performed to train the animal on the sensory consequence of a motor command and its timing. As introduced in Fig. 1E, four trial types were presented. The most common trial type Predicted presented the stimulus at the learned delay. The second class, Shifted, a test trial, was rare and presented the stimulus at a different delay than Predicted. A third class omitted the presentation of the sensory consequence (tagged ‘Omitted’, another test trial), and finally, Non-predicted, the fourth type of test trial, was presented at times when no motor activity in 7N was detected (i.e. 7N activity was below the baseline). The trial types [Predicted, Shifted, Omitted, Non-Predicted] were presented in pseudo-random order throughout the experiment with a frequency of [77, 7.7, 7.7, 7.7]%. Details for stimulus delays used in each mouse/session are given in table 1. For the experiments shown in figures 3 and 4 the delay of Predicted was always 100 ms.

**Figure 4:**
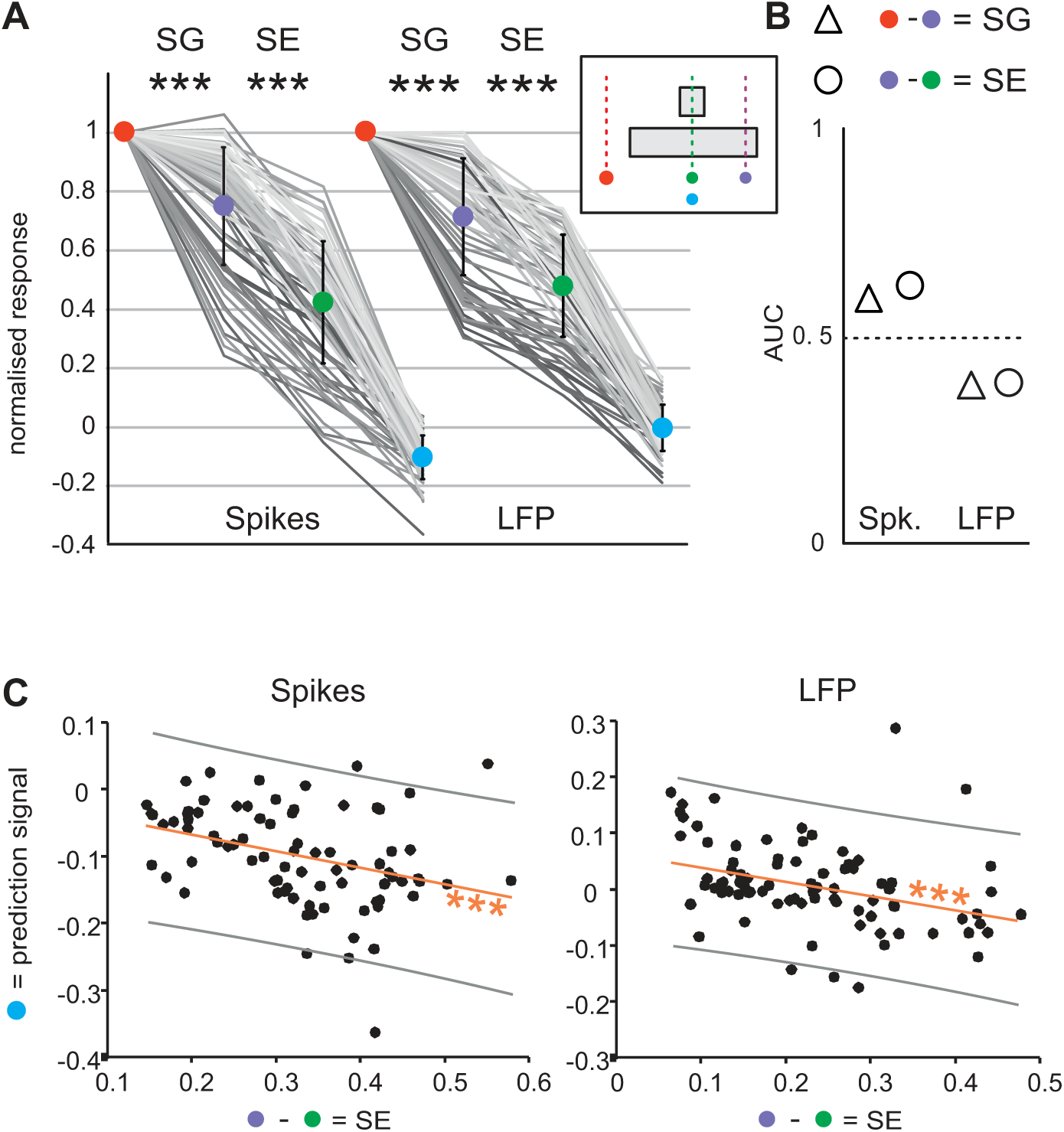
Population data of responses to different trial types. **A:** All spike and LFP responses (12 mice, for spikes, n=80 delay conditions, 176 sessions; 10 mice for LFPs, n=83 delay conditions, 157 sessions) normalized to the response to Non-predicted (red). Each line connects data from the four trial types obtained in one delay condition (the shade of gray is used to optically differentiate the lines). The inset repeats the working hypothesis (from Fig. 1E) for reference. **B:** Average AUC effect sizes of the data shown in A. **C:** Scatter plot of SE vs. the prediction signal as measured in Omitted. Orange line: regression line. Gray lines: 95% confidence interval. **Conventions for Figs. 4-10**: Significance levels: n.s.: p>0.05; *: p<0.05; **: p<0.01; ***: p<0.001; Trial types: Non-predicted = red, Shifted = violet, Predicted = green, Omitted = blue. SG is the difference between response to Non-predicted and Shifted (red minus violet); SE is the difference between response to Shifted and Predicted (violet minus green).

Figure 3D exemplifies raw 7N and S1 activity with respect to the sensory consequence (or its omission) in the four trial classes. The single trial LFP responses (colored) shown at the bottom of the panel were chosen to exemplify the consistent result as obtained in all mice reported in this study: Non-predicted evoked the largest response amplitude, followed by Shifted, Predicted and finally Omitted, which regularly gave no LFP response or a positive one. Figures 3E-G exemplify typical S1 responses (averaged across trials obtained in one session) on a much smaller time scale. As expected from previous work ^15^, S1 spiking was elevated during whisking (Fig. 3E top). After subtraction of pre-stimulus activity, we consistently observed the ranking of response magnitudes (already introduced in figure 3D) with the highest response in Non-predicted, followed by those in Shifted, then Predicted, and finally a slight spike suppression in Omitted (Fig. 3E bottom). This ranking was expected from the hypothesis depicted in figure 1EF. It states that the attenuation from the response in Non-predicted to the response in Shifted relates to the effect of SG, and the attenuation from Shifted to Predicted reflects SE (Fig. 1F). Figure 3F demonstrates two example sessions (from another animal), in which the difference in delay between predicted and shifted stimuli was varied. Decreasing the delay difference of Shifted and Predicted to 10 ms led to the coalescence of the two respective responses (a result that will be analyzed in more detail in a later paragraph). In the framework of our hypothesis this suggests that the temporal precision of SE did not allow to differentiate between SG and SE at these small delay differences anymore. Finally, figure 3G demonstrates the equivalence of effects obtained with spike rate vs. LFP recordings (note the typical negative LFP response corresponding to neuronal excitation).

The specific ranking of tactile S1 activity in response to the four stimuli (trial types) was consistently found in all 12 animals investigated. Figure 4A plots results from n=80 different *delay conditions* (the delay condition sample encompasses data points calculated across several sessions from one electrode in one animal, with Predicted at latency 100 ms and Shifted at a fixed value latency; see table 1). Results from spike and LFP data recorded from the same electrode in one animal are shown and are connected by a gray line (one gray line depicts one sample; the shadows of gray allow to follow individual lines, they were chosen purely for graphical reasons). Normalizing the responses to Non-predicted (and thus inverting the sign of LFP responses) revealed that the ranking of responses to the different trial classes (red-violet-green-blue) was observed in the vast majority of spike and LFP recordings. The strength of attenuation assigned by our hypothesis to SG (Non-predicted minus Shifted responses, i.e. red minus violet) as well as SE (Shifted minus Predicted responses, i.e. violet minus green) were found to be highly significant when tested across our sample of delay conditions (t-test, n=80 [spike rate], n=83 [LFP], p<0.001). To apply a more conservative measure, we calculated the t-tests as well based on *electrodes* (the electrode sample’s sole difference to delay conditions is that we averaged across data obtained with different latencies of Shifted) and got confirmatory results (t-test, n=42 [spikes], n=40 [LFP], p<0.001). It is noteworthy that a full attenuation of the sensory signal (i.e. to normalized response 0) by SG and SE, as depicted in the upper panel of figure 1F, was realized only in a few recordings, which may well be a technical limitation of our approach, as detecting the onset of the motor activity in 7N is prone to certain imprecision (Fig. 2G). As expected the AUC effect sizes were on average larger than 0.5 for spike recordings indicating an elevation of firing rates, and smaller than 0.5 for attenuated negative potential responses, typically found in LFP-recordings (Fig. 4B).

Omitted trials (blue) are of special interest, as one would expect them to reveal the predictive signal of SE in the absence of the learned stimulus ^10^. In fact, spike rates in Omitted showed a small but significant deviation from zero (Fig. 4A; one sample t-test, n=80 delay conditions, p<0.0001; one sample t-test, mean AUC 0.45; alternative t-test using the electrode sample: n=42, p<0.0001). However, LFP responses did not appear to be significantly different from zero (one sample t-test, delay conditions: n=83, p=0.37, mean AUC 0.49; electrodes: n=40, p=0.29). To test if this small effect with inconsistent statistical significance contains prediction value for the strength of SE-type attenuation (difference of data marked by violet and green symbols in Fig. 4A), we calculated the correlation of peak of suppression in Omitted trials with the peak response in SE in each of the 80 (spikes) and 83 (LFP) recordings (Fig. 4C). We hypothesized that stronger SE-type attenuation will be accompanied with stronger suppression in Omitted trials. This expectation was met as Omitted trial peak activity was inversely and significantly correlated with SE (r=0.25, p<0.002 for both spike and LFP data). This result argues in favor of the existence of a predictive signal for SE in barrel cortex.

Next, we analyzed the data from 5 mice obtained using across-layer recordings based on silicon arrays (Fig. 5A). Layering information was extracted from the CSD data based on the well corroborated fact that the CSD signal at a fixed latency of 5-10 ms after a pulse-like whisker deflection is known to show two sinks, one centered on L4, the other on the border of L5 and L6 (=L5/6) ^25,26^. We did not find any depth profile of the specific ranking of stimulus types. Rather, almost identical rankings were consistently observed throughout cortical depth in all five animals (Fig. 5B, depth: [-200, –100,…, +500 µm]; n (=delay conditions) for spikes: [3,6,9,9,8,8,8,5]; n for LFP [3,7,9,9,8,8,8,4]). Effect sizes of presumptive SG attenuations varied between 0.55 and 0.6 in spike data and 0.39 and 0.47 in LFPs; and SE attenuations varied between 0.60 and 0.65 in spike data and 0.33 and 0.45 in LFPs (Fig. 5C).

**Figure 5:**
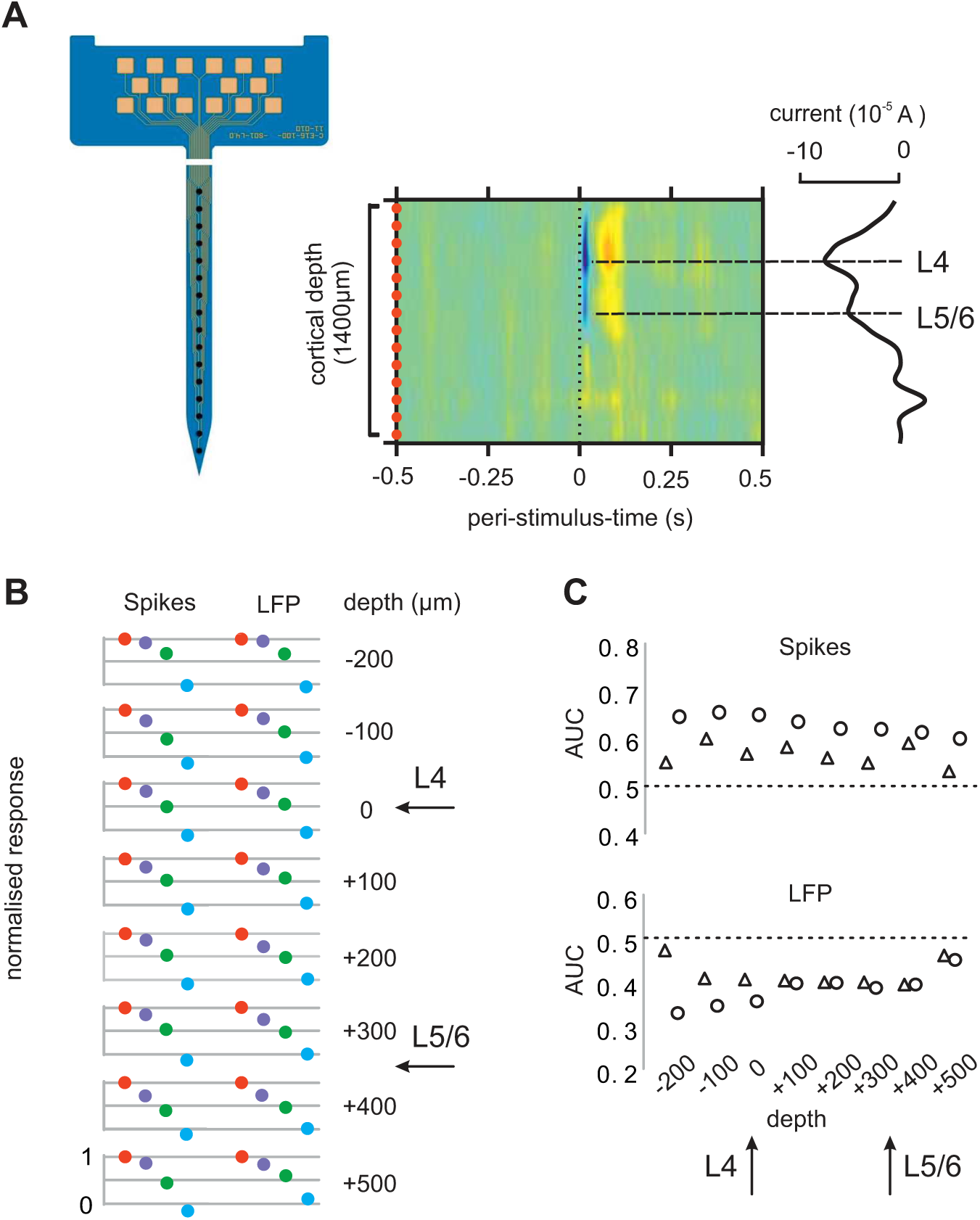
SE and SG across barrel cortex layers. **A:** Left: Silicon array with 16 electrodes, implanted across the depth of S1. Center: CSD analysis (blue: sinks, red: sources). Right: At short latency after the stimulus two sinks were observed. The two negative peaks were taken as depth marks L4 and border of L5/6 ^25^. These depth marks were transferred to panels B and C. **B:** Response to trial types through cortical depth (average of 5 mice). **C:** Average AUC effect sizes of the data shown in B. **Conventions** as in Fig. 4.

To quantify the temporal resolution of SE attenuation, we varied the delay of the stimulus in Shifted trials with respect to Predicted (Fig. 6A). We trained 5 mice on a delay of Predicted and varied the Shifted from 100 ms to 50 ms and 10 ms difference from predicted delay. The specific ranking of responses was observed consistently and significantly with a delay difference between Predicted and Shifted of 100 ms and 50 ms (t-test, n=16 for spikes and 17 for LFPs, p<0.001), but broke down using 10 ms difference (i.e. attenuation of the two trial types was not discernable anymore; t-test, n=16 (delay conditions) for spikes, p=0.663 and n=17 for LFPs, p=0.09) (Fig. 6B). Correspondingly, the effect size of SE obtained with delays larger than 10 ms indicated clear SE-like attenuation, while at 10 ms random performance was reached across the depth of S1 (AUC close to 0.5; Fig. 6D).

**Figure 6:**
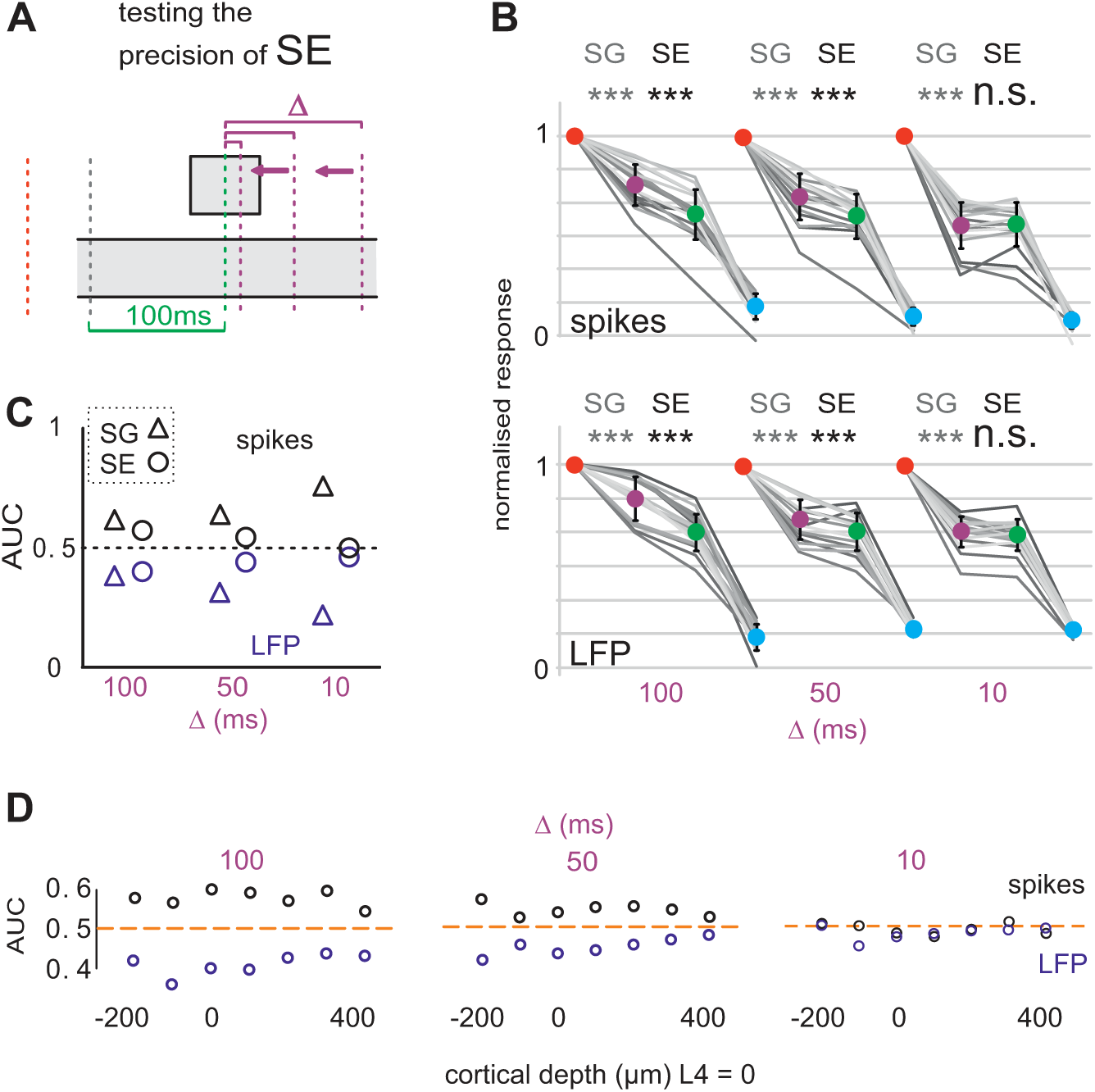
Precision of SE. **A:** Experimental approach. The latency of the Shifted stimulus was set closer and closer to the one of Predicted stimulus. The difference latency Δ was 100 ms, 50 ms, and 10 ms (violet, presented in blocks of subsequent sessions). The gray line marks movement onset. **B:** Population spike and LFP mean responses normalized to the response to Non-predicted (red). At Δ=50 ms and 100 ms, SE (black) was highly significant. With Δ=10 ms SE-like attenuation vanished. Significance of SG (gray), was unaffected, as expected (data from 5 mice). **C:** Average AUC effect sizes of the data shown in B. **D:** Average AUC effect sizes at the same Δ across layers (2 delay conditions from 2 mice). **Conventions** as in Fig. 4.

We next tested how late after onset of the motor command the SE-attenuation could be observed. To this end, we trained mice at Predicted delay of 0ms (4 mice), 200 ms (6 mice) and 400 ms (4 mice; Fig. 7). We observed in the distribution of responses from all electrodes a diminished attenuation that presumptively can be attributed to SE at 200 ms delay as compared to 0 delay (Fig. 7B). Nevertheless, both attenuations were highly significant (delay 0 ms: t-test, spikes: n (delay conditions) =9, LFPs: n=11, p<0.0001; delay 200 ms: n=23 spikes, n=30 LFPs, p<0.0001). Finally, the SE-attenuation vanished when increasing Predicted at latency 400 ms (t-test, spikes: n=20, LFPs: n=24, p>0.05), while, importantly, the attenuation attributable to SG stayed high. Figure 7C shows the same result plotting the average effect sizes, while figure 7D demonstrates average effect sizes of SE across layers in 4 mice. We next tested how fast an attenuation for a changed stimulus delay can be learned (Fig. 8). In additional experiments the delay of Predicted was trained at 100 ms and then switched from one session to the next to 200 ms (while Shifted was kept the same across the switch). We compared the SE-attenuation in the session before a switch, and after the switch in 4 mice. We found that the animals already after the first post-switch session reached an attenuation that was a little smaller as compared to the one expressed pre-switch, but was highly significant for spike rates as well as for LFPs (t-test, n=15 spikes, n=14 LFPs, p< 0.001) (Fig. 8A). Accordingly, effect size for SE-attenuation was separate from chance but slightly reduced in the post-switch session (Fig. 8B). The animals, therefore, successfully learned a new Predicted stimulus delay within a single behavioral session.

**Figure 7:**
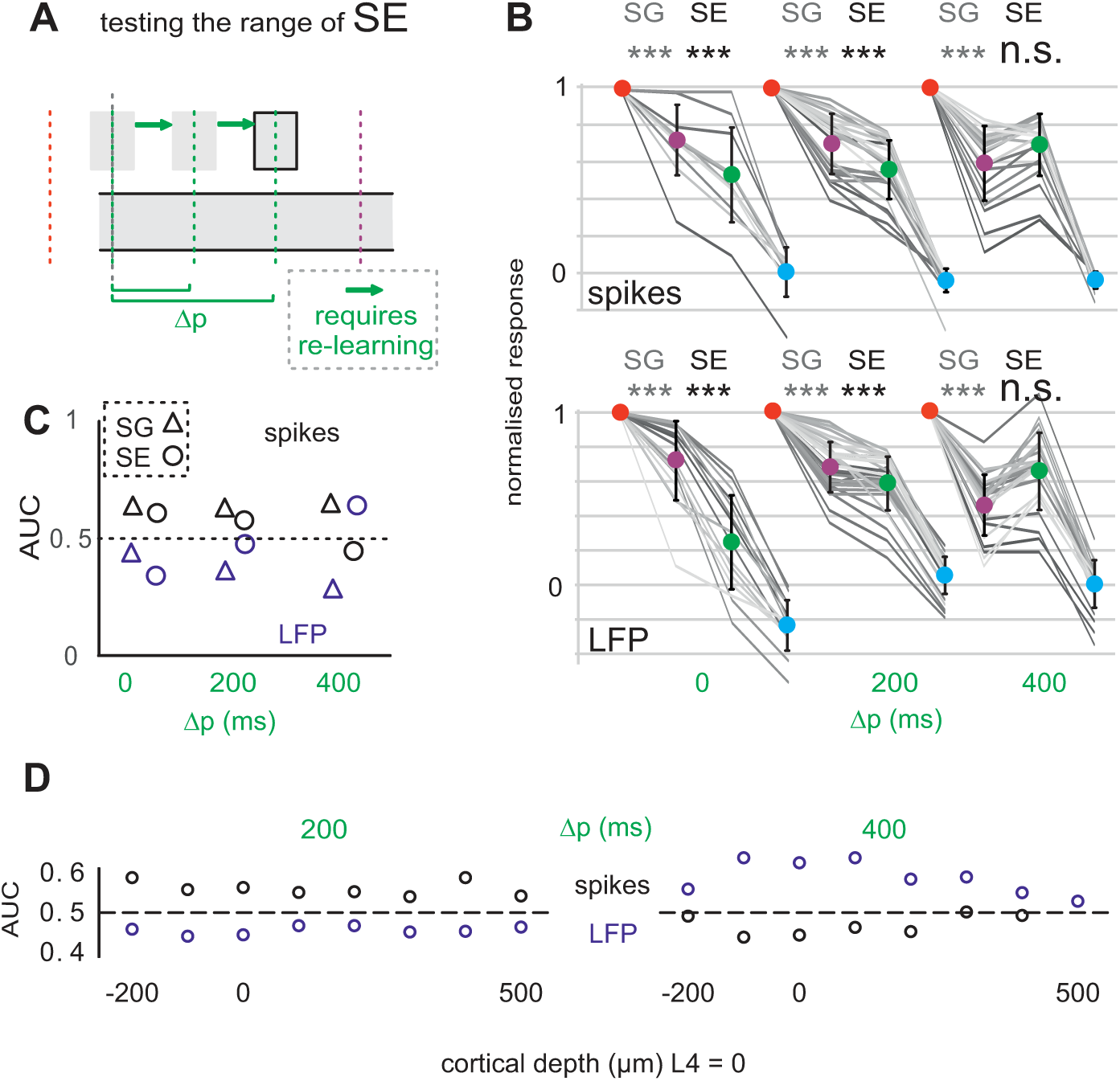
Range of SE. **A:** Experimental approach. The latency of the Predicted stimulus was moved across the post-trigger time. New latencies were presented in subsequent blocks of sessions. Whenever a change in latency was presented it had to be learned by the animal. The latencies of Predicted from onset of motor command were Δp = 0 ms (4 mice), Δp = 200 ms (6 mice), and Δp = 400 ms (4 mice). The gray line marks movement onset. **B:** Population spike and LFP mean responses normalized to the response of Non-predicted (red). At Δp = 0 and Δp = 200 ms SE was highly significant (black). At Δp°=°400 ms the SE effect vanished. Significance of SG (gray), was unaffected, as expected. **C:** Average AUC effect sizes of the data shown in B. **D:** Average AUC effect sizes at the same Δp across layers (3 delay conditions from 3 mice). **Conventions** as in Fig. 4.

**Figure 8:**
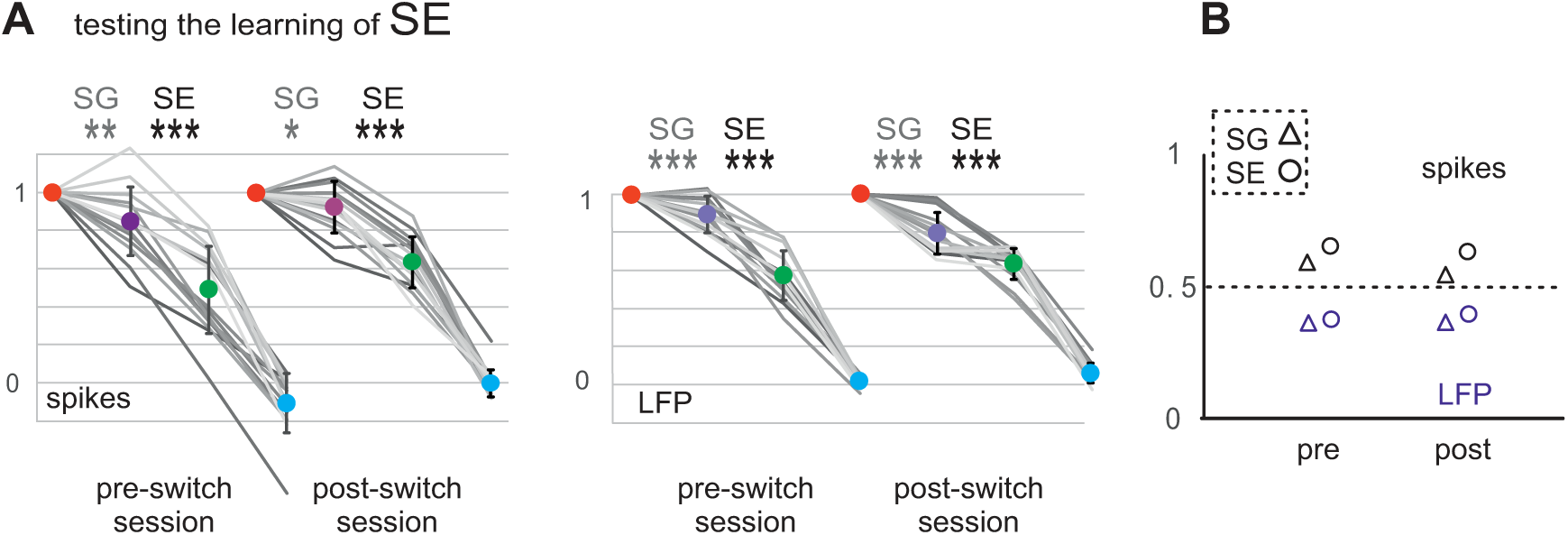
Learning a new Predicted latency is faster than one training session. **A:** Population spike and LFP mean responses from a session before the change in latency (pre-switch) and after (post-switch) normalized to the response of Non-predicted (red) (data from 4 mice). SE was highly significant in the pre-switch session, but also already in the first post-switch session. Expectedly, the significance of SG (gray), was stable as well. **B:** Average AUC effect sizes of the data shown in A. **Conventions** as in Fig. 4.

As mentioned already, SG-like attenuation was visible at delays, at which SE-like attenuation already had vanished. To find out how long after the onset of the motor command SG-like attenuation can be activated, we fixed Predicted to a delay of 100 ms, while Shifted changed, in a session-wise fashion, from 500 ms (6 mice) to 700 ms or l000ms (4 mice). Figure 9A shows population data of the sessions with a delay of Shifted of 500 ms and 700 ms and larger. As expected from our results shown in figures 3 and 4, SE-attenuation stayed highly significant, independent on Shifted delays. Our focus, however, was on the SG-attenuation, which was significantly present in sessions presenting Shifted at a delay of 500 ms, and only vanished when the shifted delay moved to 700 and 1000 ms (n= 21[delay conditions], p>0.05 spikes; n=23, p<0.05 opposite direction LFPs). Therefore, the time point, after which SE (200-400 ms) and SG (500-700 ms) cannot be evoked anymore is different (cf. Fig. 7 and 9), and thus lends another argument for the separation of the two mechanisms. The same data analyzed to yield effect sizes are plotted in figure 9C, and split according to recordings from electrodes located in different cortical depths (Fig. 9D).

**Figure 9:**
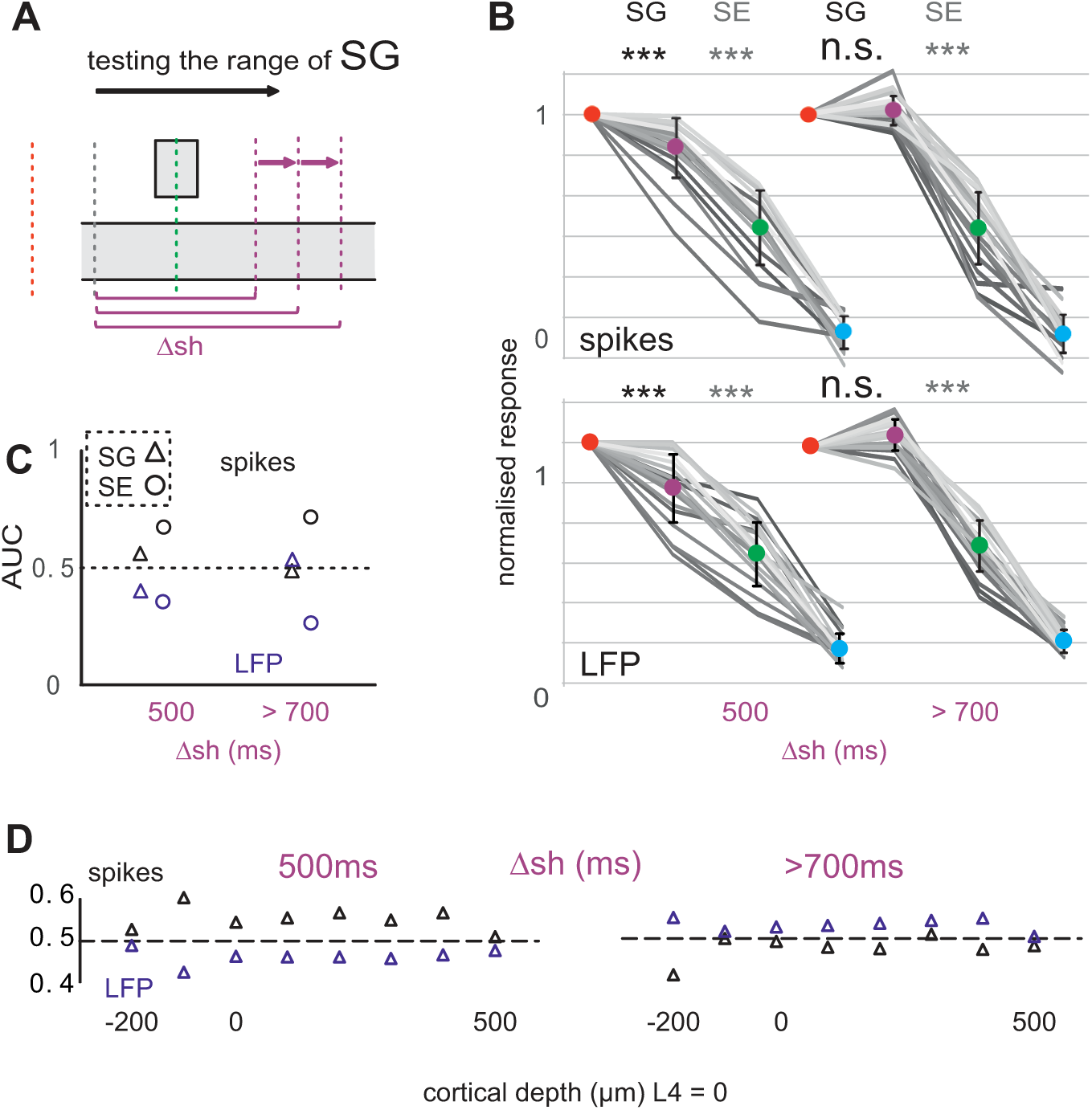
Range of SG. **A:** Experimental approach. The latency of the Shifted stimulus was moved across the post-trigger time. New latencies were presented in subsequent blocks of sessions. The latencies Δsh = 500 ms (6 mice), Δsh = 700 ms (2 mice) and Δsh = 1000 ms (2 mice) were presented. The gray line marks movement onset. **B:** Population spike and LFP mean responses normalized to the response of Non-predicted (red). At Δsh = 500ms, the SG-like attenuation was significant (black), but vanished with Δsh = 700 ms or higher. Expectedly, the significance of SE (gray) was observed to be stable under these manipulations. **C:** Average AUC effect sizes of the data shown in B. **D:** Average AUC effect sizes at the same Δsh across layers (5 delay conditions from 5 mice). **Conventions** as in Fig. 4.

### Closed loop experiments

During open loop experiments whiskers were immobilized, impeding a behavioral read-out of the engagement of the motor system. To control for the possibility that mice changed task strategies in unknown ways after lesion of the facial nerve, we acquired data from four additional mice that were trained in closed loop control (Fig. 10AB), keeping the whisker movement as a behavioral read-out throughout the experiment. The sensory consequence was presented via electrical stimulation of principal trigeminal nucleus (Pr5), because the whisker movement would impede precise delivery of a sensory consequence in form of a whisker flick as done before. All results obtained using the closed loop method mirrored the results as observed under open loop conditions: A Pr5 micro-stimulation pulse evoked spike and LFP responses in S1 at short latency of 3-5 ms ranking the trial types as observed before (red-violet-green-blue; Fig. 10C). The population data (Fig. 10D) demonstrates the separation of SE and SG types of attenuation, suggesting that results in open and closed loop experiments were similar. As in open loop experiments (cf. Fig. 4), we used a delay of 100 ms for Predicted trials but recorded sessions with different Shifted. As before, we calculated the statistics using the delay conditions sample as well as the electrodes sample.

**Figure 10:**
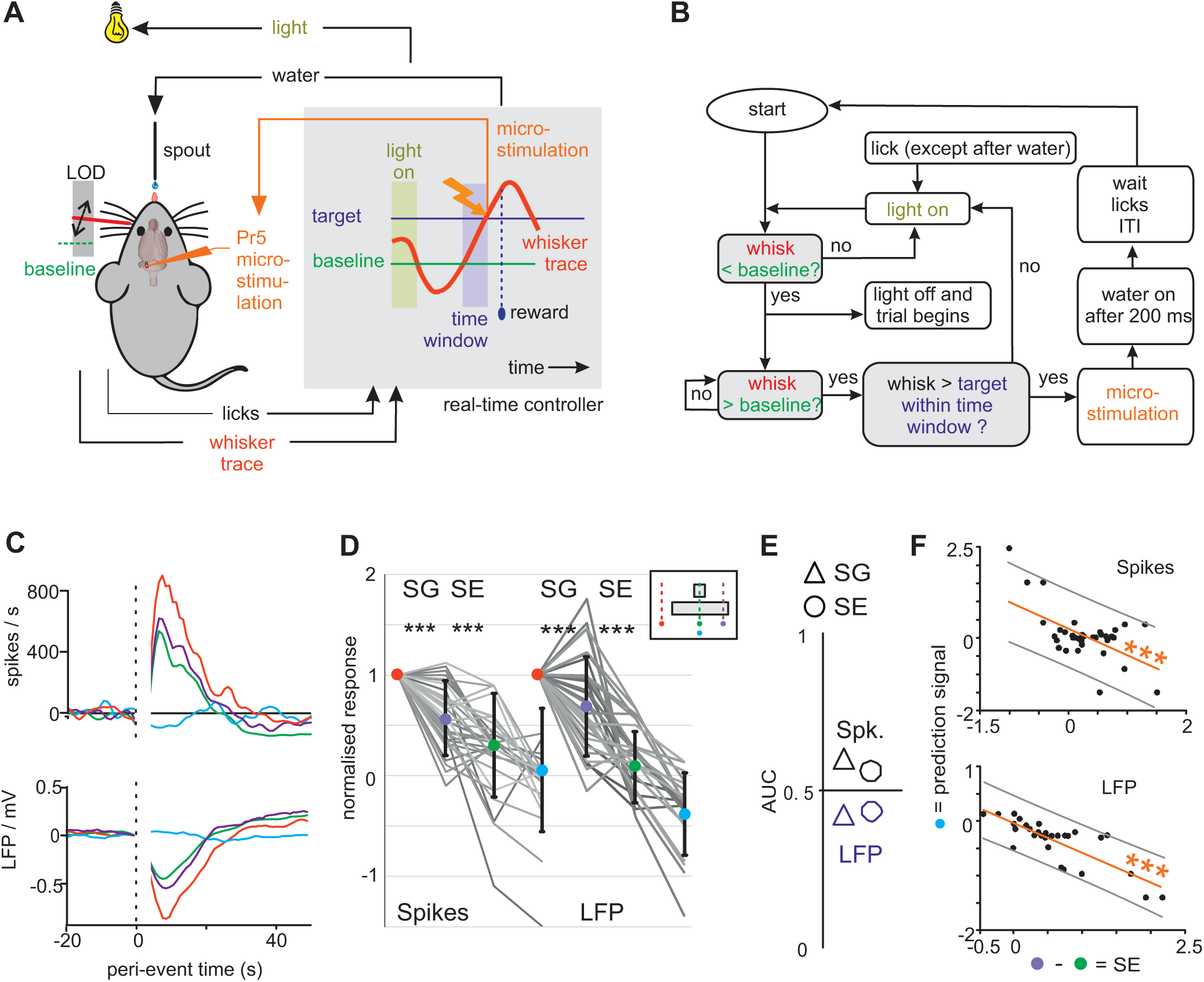
Closed Loop Experiment: Training of whisker reach with tactile consequence. **A:** Upon a successful whisker reach movement (see Fig. 2), the real-time controller generates a trigger signal that is converted into a tactile consequence via a microstimulation pulse in principal trigeminal nucleus (Pr5). **B:** Flow diagram of the state engine realizing automatic experimental control. **C:** Example responses from one recording site. Top: Spike rates (multi-unit recordings). Bottom: LFP responses. **D:** Population data (4 mice, delay conditions: n=46 for spikes and n=30 for LFPs). Each line connects responses to the four stimulus types for one delay condition (Predicted delay is always 100 ms) (the shade of gray is used to visually differentiate the lines). The responses are normalized to the response to Non-predicted (red). The inset repeats the working hypothesis for reference (cf. Fig. 1E). **E:** Effect sizes. **F:** Correlation of SE with the prediction signal (response to Omitted). **Conventions** as in Fig. 4.

The results were similar in the two cases, and they were comparable to the ones obtained using the open loop approach. The t-test using delay conditions as samples gave: SG: t-test, n=46 [spikes], p<0.001, n=30 [LFP], p<0.01; SE: t-test, n=46 [spikes], p<0.001, n=30 [LFP], p<0.001. The one across electrodes after averaging across different shifted conditions gave: SG: t-test, n=11 [spikes], p<0.001, n=8 [LFP], p<0.01; SE: t-test, n=11 [spikes], n=8 [LFP], p<0.05. The average effects sizes were above and below 0.5 for spikes and LFPs respectively (Fig. 10E), and the predictions signal recorded with Omitted trials was significantly correlated to SE (p<0.001; Fig. 10F), corroborating the findings seen in the open loop experiments.

## Discussion

Imposing experimental sensory consequences using open and closed loop sensorimotor experiments based on a whisker reach task in mice, we found consistent evidence for two separate neuronal mechanisms, SE and SG, both of which led to an attenuation of the tactile responses in S1. Our arguments for the separability of the two mechanisms are the following: First, they show different relation (and precision) to the experimental sensory consequence. SE was found to be active solely within <50 ms around a point-like sensory consequence. Second, the time window after the onset of a motor command / movement in which they can be evoked differs. SE showed characteristics of the classical expectation from the reafference principle ^10^: it attenuated the tactile signal only at the predicted time of a sensory consequence. The function of SG remains obscure. Clearly independent from the sensory consequence, it was loosely related to the movement. An important speculation to be tested further is that its long active interval after the motor command / movement onset may point to a function beyond details of motor control, like attentional functions and/or reward prediction.

### Open vs. closed loop

State estimation can be tested either with movement perturbation ^7,8,27^ or by adding experimental sensory consequences. The latter can be done employing the open loop ^10^ or closed loop approach ^11,28^, i.e. preventing the natural sensory feedback (open) or leaving it in place (closed). As our aim in this study was to delineate two predictive systems our choice was to add different sensory consequences as this approach let us test the additional existence of SG in straightforward manner (cf. Fig. 1E).

There are pros and cons for open vs closed loop approaches. Testing predictive signals in open loop condition abolishes relevant sensory inputs and therefore argues strongly in favor of the contribution of a centrally generated predictive signal, likely originating from an internal model. A drawback is the possibility that after preventing the movement (and its read-out) the behavior may change unbeknownst to the experimenter. In closed loop, the disadvantage is that the movement and natural feedback (the reafference) remains intact, which at least in principle allows for the possibility that the observed sensory attenuation is based exclusively on sensory feedback without a necessary contribution of an internal model. In view of these drawbacks, we decided to run both, open and closed loop experiments to test if the obtainable results confirm and reinforce each other. In fact, the result of both experiments was identical within the confines of our experimental precision. We, thus, can exclude with reasonable confidence, that the attenuations are not due to changes in behavioral strategy (from the closed loop experiments). Second, we strengthened the argument for the idea that the attenuations very likely originate from a central signal (from the open loop experiments).

To the best of our knowledge our results are the first to demonstrate SE in the whisker system, and they are the first to show SE using an open loop approach in a mammalian animal model. The suggestion on SE’s dependence on central signals strengthens the notion that state estimation uses an internal model, most likely located in the cerebellum ^6^. In the case of SG our result of its existence and dependency on central signals is confirmatory, as our previous work has shown that it survives even a cut of tactile primary afferents in the infraorbital nerve ^15^.

### Separability and function of SE and SG

Our arguments for the presence of two separable types of sensory attenuation rest, firstly, on the dependence/independence from a learned sensory consequence of movement, and secondly, on the intervals after movement / motor command onset, in which SE and SG could be evoked. Most of the work done to study SE and SG in rodents and primates falls apart into groups of studies, studying either one or the other, using different experimental approaches. We therefore start the discussion with one recent work that, like our present study, has attempted to study two predictive systems in the same experiment ^16^. We think that the results of the prior study, based solely on human psychophysics, has revealed some behavioral aspects of the phenomena we have described here using neuronal recordings. Having participants determine touch intensity on their left hand evoked by movement of the right one, with and without movement of the left one, Kilteni and Ehrsson found two separable suppressive perceptual mechanisms, which they called ‘attenuation’ and ‘gating’. ‘Attenuation’, was a diminished tactile percept evoked in the left hand that in predictable ways was tapped by a movement of the other. There is a rich literature on force matching tasks, reporting that this attenuation depends on the presence of a movement command in a temporal precise way, and on predictive signals originating from the cerebellum ^4,18,29,30^. Tight temporal relationship to the motor command argues in favor of the possibility that the ‘attenuation’ observed by them corresponds to what we call SE. ‘Gating’, their second phenomenon, independently suppressed perception of a tap presented at the left hand at unpredicted times during the movement of the left hand. This resembles the characteristic of SG as reported here.

So far, no direct experimental facts about neuronal separation of SE-and SG-like attenuation in primates is available. However, whenever, SE-or SG–like phenomena were studied separately in humans ^4,18^ or primates ^31^ combined with any assessment of neuronal activity, S1, S2 or surrounding parietal cortex has been found to be active, furthering our notion that SE and SG phenomena, as consistently found here in rodent S1, may have a close correspondence in the primate brain.

Previous works in the whisker system have studied sensory attenuation during movement, but never with the temporal precision and link to the onset of a movement as done here ^13,15,32–34^. Therefore, we hold it likely that in the whisker system, so far, exclusively the effects of the SG predictive system have been reported. While our present approach aimed at delineating SG from processes akin to reafference principle and state estimation, our experiments provide little additional insight toward the behavioral function of SG. We note that SG-like attention firstly has been found to dependent on parietal cortical structures in rodents and is projected down the ascending tactile pathway ^13^. Secondly as mentioned already, the fact that SG survives blockade of peripheral efferent and afferent signal flow (this study and ^15^), excludes with high certainty the possibility that it relies on any movement dependent sensory signals. Lastly, studying arm movements in primates, it has been found to be active around 150 ms before the initialization of the movement ^31^. These findings strengthen the notion that SG indeed is based on an internal (and therefore potentially predictive) signal and they point to a much looser relation of SG to details of movement command or trajectory as compared to SE. We wish to point out that in many of the previous studies describing SG-like attenuation in rodents or primates, complex motor or perceptional tasks had to be performed with the aim to obtain a reward, which opens the possibility that SG-type of prediction may well be situated on a hierarchically higher level, as required for instance for attentional processes or reward prediction.

The characteristics of SE, the dependence on a predicted sensory consequence as well as its time range, matches those reported from cerebellar-like structures in fish ^10^ and mice ^11^. In these previous works, artificial sensory consequences (electric shocks in the fish tank ^10^, as well as acoustic emissions brought about by licking movement at water dispensers ^11^) were predicted and attenuated. We consider the similarity of experiment and the finding of specific attenuation of tactile flow a strong suggestion that within the tactile whisker-related system, SE in S1 is the reflection of a neuronal circuit that is involved in comparing an internal predictive signal and sensory feedback, i.e. the very process of state estimation ^6^. The persistence of SE in open loop is consistent with the involvement of an internal model. Paw reach perturbation experiments in mice have pointed to S1 as the seat of the comparator of internal model predictions and sensory feedback ^12^. Our findings in the whisker system are, in principle, consistent with this idea, as we found SE-like attenuation, as well as predictive signals (in Predicted and Omitted trials respectively) in S1. However, we note that there exist multiple places in the brain that have the potential to integrate tactile signals with cerebellar predictions, amongst them the cerebellar nuclei (assuming that predictions are generated by the cerebellar cortex), thalamic stations mediating cerebellar signals to the neocortex, as well as the ascending tactile pathway in the brainstem. All of these structures may readily have access to the inferior olive ^35–37^, the structure thought to generate the error signals needed to update the cerebellar cortical internal model ^38,39^. Before a final verdict on the location and mechanism of the comparison of sensory feedback with internal predictions can be issued, these stations have to be included into the picture as well.

## Methods

### Animals

All procedures were approved by the local authorities as required by German law. We used male mice of the strain C57BL/6N, aged 13 weeks, obtained from Charles River Laboratory (Germany). The animals were kept at an inverted light cycle (12h/12h, dark during the day, light during night) at 25° room temperature and 55% humidity. Behavioral experiments were performed in the same room during the day, the active phase of the animals. Behavioral procedures of water control, habituation to head fixation, and behavioral training were done as described in a technical article ^40^.

### Electrode implantation into 7N and Pr5

Mice were implanted with a single tungsten electrode (1-3 MΩ) in 7N in the area that corresponds to whisker protraction. Anesthesia was induced by intraperitoneal (i.p.) injection of 3K (fentanyl 0.5 mg/kg, midazolam 12.5 mg/kg, and fluanisone 25 mg/kg, i.p.). Supplements of 33% of the initial dose were given approximately every 50 mins repeatedly checking the response to tail/tow pinch. The body temperature, monitored by a rectal probe, was held higher than 35°C by a warming pad. The dorsal head was shaved, and the mouse was mounted in a stereotaxic frame. The skin above the dorsal skull was disinfected, incised, connective tissue and periost locally anesthetized with Lidocaine (1 %) and bluntly stripped to the sides. The dry bone was prepared using an light-polymerized adhesive (Optibond, Kerr, Germany), which later made the contact with dental cement. A craniotomy over 7N (diameter maximally 1.5 mm at 5.5 mm rostrocaudal from bregma and 1.8 mediolateral from midline) was drilled. (spherical drill bit at 5000 rpm). The brainstem starting at of depth of 4.8 mm from the cerebellar surface was then mapped using microstimulation (a pulse train of biphasic rectangular pulses of 5-10μA current [negative current first]) while monitoring movement of the whiskers through a binocular. After locating an area that gave rise to brisk whisker protractions, the surface of the brain was covered by Kwik-Seal (World Precision Instruments, USA) and the electrode and its plug embedded by dental cement (Tetric EvoFlow, Ivoclar, Germany). This electrode served to record the whisker motor command in the later experiment. The wound was then sutured to adapt closely around the headcap. Analgesia was induced by carprofen (5 mg/kg, i.sc.) followed by the 3K antidote (naloxone 1.20 mg/kg, flumazenil 0.50 mg/kg, and atipamezole 2.50 mg/kg, i.p.), upon which anesthesia was reversed within a few minutes. Post-surgery care included the continuation of carprofen injections, the administration of warmth and, if needed, electrolytes. Twelve animals received 7N electrode implantation.

Four mice were prepared for control experiments in ‘closed loop’. For this purpose, no 7N electrode was required. Instead these mice received a single microelectrode implantation in Pr5 in the first surgery (small craniotomy centered at 5mm rostro-caudal from bregma and 0.8 mm mediolateral from the midline, starting at a depth of 4mm from the brain surface). Pr5 was identified by recording neural response to whisker deflections using a hand-held cotton swab. The electrode was implanted at a site that gave rise to spike responses of deflections of whisker C2. This electrode served to apply microstimulation pulses into Pr5 as a movement-triggered sensory consequence.

### Electrode implantation into barrel cortex and preparation of head post

After complete recovery from the first, a second surgery (anesthesia as described above) was performed. Optical imaging exactly done as before ^41^ was used to locate barrel column C2. Guided by markings at 0.5mm-2.5mm rostrocaudal from bregma and 2.5-4mm mediolateral from midline, applied in the first surgery, the skull was thinned carefully using a drill bit at 1000 rpm. The electrode array was lowered in steps of 1µm into the cortex and fixed at a depth of 300-350 µm using dental cement, after covering the craniotomy with Kwik seal. A head-post (M1 stainless steel screw, head-down) was placed on the back of the skull and embedded within the dental cement. The skin was sutured at the ends of the incision until a close adaptation, free of mechanical stress, of skin and dental cements was achieved. Post-operative treatment was the same as above.

### Electrophysiology

Custom-made platinum/tungsten-in-glas electrodes were pulled and grounded in-house and used either as single electrodes (for 7N and Pr5 implantations) or fashioned into movable 2×2 arrays (for barrel cortex implantation, Haiss et al., 2010). The length of the electrodes was 6-7 mm for 7N and Pr5 and 2mm for barrel cortex implantations. Silicon arrays, single shank containing 16 electrodes of impedance around 1 MΩ, were also used in some animals to implant in barrel cortex (length of the shank 1500μm, distance between electrodes 100μm) (E16-100-S1-L6 NT, Atlas Neuroengineering bvba, Belgium). Silver wire electrodes were implanted onto the dura mater over the cerebellum and prefrontal cortex as ground and reference electrodes, respectively. The wire end was fashioned into a ball, to minimize damage of dura and brain.

From the total sample of 16 mice all received silver ball electrodes; 7 received 7N wire and barrel cortex wire array electrodes; 5 received a 7N wire electrode, and barrel cortex silicon array electrodes; 4 received a Pr5 wire electrode, and barrel cortex wire array electrodes.

Multielectrode extracellular recordings were done against a silver-ball reference (multichannel systems, Reutlingen, Germany, sampling rate: 40 kHz). The local field potential (LFP) was obtained by down sampling the data to 2 kHz and band pass filtering of the voltage recordings (Butterworth, edge frequencies 1 and 200 Hz; the filter order was assigned to reach passband ripple amplitude of less than 0.5 dB and stopband attenuation exceeding 30 dB). Spike recordings were obtained by setting the bandpass edge frequencies to 500 to 5000 Hz. The resulting voltage traces were thresholded to obtain multi unit spike time stamps. For closed loop experiments, the artifact produced by electrical stimulation was removed. This was achieved by subtracting the median response across trials from the signal followed by blanking the window [0, 4] ms after stimulation onset.

### Training whisker reach

After complete recovery from all surgical interventions, and habituation to head-fixation ^40^, the mice were trained on a whisker reach task with the same equipment and exactly following the strategy as reported before ^22^. After the behavioral training, mice consistently and precisely performed reaching movements (no touch) of their left whisker, starting from a predefined baseline location or behind (located just rostral to their whisker resting position), and surpassing a threshold location more rostral. They were trained to consistently reach across an amplitude of 45° within a time window of 50 ms. This training took typically 2-3 weeks.

### Open and closed loop experiments

In the 12 mice investigated in open loop conditions, the facial nerve was cut in a short surgical intervention (15 min). Anesthesia and postoperative care were as described above. The ipsilateral cheek area was shaved, and a vertical incision was made. Muscles were retracted to locate the buccal and marginal branches of the facial nerve. A 1 mm long section was removed from both buccal and marginal branches to reduce the probability of nerve regeneration which led to immobility of the whiskers on the left side. The neural activity was recorded and analyzed by a brain-machine interface based on a state engine controlling the experiment (Simulink, Natick, MA, USA, the state engine ran on a real-time PC at a framerate of 10 kHz). The motor command recorded in 7N was then used to trigger the artificial sensory consequence, a sinusoidal whisker deflection (from one negative extreme to the next, amplitude: 10°; duration: 10 ms). To identify 7N motor commands, the state engine rectified and integrated the 7N spike recording at each time step. It used two parameters to decide whether in a past window (0 to 500 ms before the actual time) the activity was below threshold A and in the present window (0 ms before the present time) it was larger than threshold B. The values of A and B had been determined by the experimenter in the sessions before the facial nerve was cut to optimize the detection of whisker movement commands from 7N activity. Once a valid motor command was detected, the above described whisker flick was triggered at the preset latency followed 200 ms later by the delivery of a drop of water via a spout in front of the mouse’ snout (Figs. 2 and 3).

In four mice closed loop experiments were performed as a control. That is, no facial nerve cut was performed, and the tracking of the intact whisker reach movements were used to inform the state engine about the state of the animal’s behavior (Fig 10 AB). The time point when the whisker crossed the threshold was used to trigger the sensory consequence at the preset delay. The water reward was triggered after passing the threshold at a delay of 200 ms. In these experiments, the sensory consequence was the presentation of a single microstimulation pulse in the C2 barrelette of Pr5 (bipolar rectangular current; amplitude: 10 µA; duration: 300 µs).

Trials were grouped into 4 classes. The first and most common trial type, called ‘Predicted’, presented the stimulus at the learned delay from motor command or movement onset (Fig. 1A). The second trial type, called ‘Shifted’, presented the sensory consequence at time point, which was still inside the interval in which the whiskers were moving after the trial onset (a few hundred ms). The third class (‘Omitted’) would omit any sensory consequence, and the fourth class, called ‘Non-predicted’, would present the sensory stimulus in absence of a motor command / movement. A behavioral session would present the four stimulus classes in a pseudo-random sequence that contained a series of blocks of 13 trials (seamlessly presented without any additional gaps, etc.), in which 10 trials were of type Predicted; one Shifted, one Omitted, and one Non-predicted.

### Data analysis

To judge the depth of silicon array recording sites within the depth of the barrel column the multisite LFP signals were transformed into a current source density map (CSD). We employed a kernel-based current source density method ^43^, which generated a two-dimensional CSD space spanning the silicon shank and peri-stimulus time (Fig. 5). The pattern of sinks and sources in barrel cortex, a few milliseconds after a brisk whisker stimulus, is well-documented ^25,26^ and allowed us to assign the location of layer 4 (L4) and the border between layers 5 and 6 (L5/6) within the 16 recordings on the silicon shank. The depth scale was established by using the peak of the sink at L4 (position zero) and the electrode spacing on the silicone shank to be able to compare data across mice.

Firing rates (number of spikes multiplied by the number of trials divided by the duration of a time bin) were calculated from the four trial types (at 1 ms duration of a time bin). The non-parametric effect size AUC, based on ROC analysis ^44^ was calculated for spike and LFP responses. Ensembles of spike counts, or LFP values across trials (obtained during the response to the stimulus) were used as input to the analysis. AUC assumes a value of 0.5 if the two response distributions are indistinguishable, and extrema of 0 or 1 if they are fully separated. We always observed AUC > 0.5 for excitatory spike responses, while the corresponding values for LFPs data were typically < 0.5 (because our analysis focused on the depth between L4 and L5/6, at which local spiking is reflected as negative LFP). Note that, while the graphs plotting AUC in this article readily show this sign conversion, the graphs depicting neuronal responses do not. Instead they plot normalized effects for better comparability (e.g. compare panels 4A and B).

## Acknowledgements

This research was supported by grants from the Deutsche Forschungsgemeinschaft (SCHW577/21-1, SCHW577/22-1). We thank Alia Benali for discussions, and Ursula Pascht and May Li Silva-Prieto for technical and organizational help.

## Author contributions statement

KG designed and conducted all experiments, analyzed data for open loop experiments and wrote the paper. RRC designed and conducted closed loop experiments, analyzed data, and wrote the paper. SC designed open loop experiments, and analyzed data. CS designed all experiments, analyzed data, and wrote the paper.

